# Microglial NF-κB Signaling Deficiency Protects Against Metabolic Disruptions Caused by Volatile Organic Compound via Modulating the Hypothalamic Transcriptome

**DOI:** 10.1101/2023.11.08.566279

**Authors:** L.K. Debarba, H.S.M. Jayarathne, L. Stilgenbauer, A.L. Terra dos Santos, L. Koshko, S. Scofield, R. Sullivan, A. Mandal, U. Klueh, M. Sadagurski

## Abstract

Prolonged exposure to benzene, a prevalent volatile organic compound (VOC), at concentrations found in smoke, triggers hyperglycemia, and inflammation in mice. Corroborating this with existing epidemiological data, we show a strong correlation between environmental benzene exposure and metabolic impairments in humans. To uncover the underlying mechanisms, we employed a controlled exposure system and continuous glucose monitoring (CGM), revealing rapid blood glucose surges and disturbances in energy homeostasis in mice. These effects were attributed to alterations in the hypothalamic transcriptome, specifically impacting insulin and immune response genes, leading to hypothalamic insulin resistance and neuroinflammation. Moreover, benzene exposure activated microglial transcription characterized by heightened expression of IKKβ/NF-κB-related genes. Remarkably, selective removal of IKKβ in immune cells or adult microglia in mice alleviated benzene-induced hypothalamic gliosis, and protected against hyperglycemia. In summary, our study uncovers a crucial pathophysiological mechanism, establishing a clear link between airborne toxicant exposure and the onset of metabolic diseases.

## Introduction

Air pollution has emerged as a significant risk factor for both obesity and Type 2 Diabetes (T2D) ^1,2^. Among the common air pollutants, volatile organic compounds (VOCs) stand out due to their widespread presence indoors and outdoors, mostly emitted during vehicle operation, fuel production, and exposure to smoke ^3–5^. However, research on the metabolic consequences of exposure to these non-occupational VOC remains limited. Nevertheless, recent epidemiological studies demonstrate correlation between environmental exposure to benzene, a prominent VOC in the air, and the risk for developing insulin resistance. Studies done with children, adolescents, and elderly adults living in polluted areas and exposed to low levels of benzene, showed association between urinary benzene metabolites and the development of insulin resistance ^6–10^. Further, benzene exposure was associated with increased cardiovascular risk in both smokers and nonsmokers ^11^. Together, these studies suggest that exposure to environmental VOC represent an underestimated risk factor for metabolic abnormalities.

Mutual relationships between metabolic abnormalities and the immune system were reported particularly in obesity, T2D, and cardiovascular disease ^12,13,14^. Accordingly, numerous studies show association between exposure to various VOCs including benzene, and increased systemic inflammation in diverse urban populations ^15–17^. In our previous study we reported that chronic exposure to benzene at levels mimicking cigarette smoking, led to insulin resistance, hyperglycemia, and increased expression of inflammatory genes associated with the IKKβ/NF-κB signaling pathway in the liver and the hypothalamus, specifically in male mice ^18^. These findings underscore the potential implications of benzene exposure in the development of metabolic disturbances, with particular relevance to the critical organs involved in metabolic regulation.

The hypothalamus receives input from a variety of hormonal, environmental and nutritional signals, which work together to regulate an organism’s glucose metabolism and energy balance ^19^. In the context of metabolic disorders like obesity and diabetes, neuroinflammatory responses characterized by dysregulated glia cells in the mediobasal hypothalamus (MBH) have become a notable hallmark ^20,21^ ^22,23^. Microglia, the central nervous system (CNS)-resident immune cells, exhibit sensitivity to stressors and can be activated by inhaled components of urban air pollution through both direct and indirect pathways ^24–26^. This stress drives microglia to release interleukin 1α (IL-1α), tumor necrosis factor α (TNF-α), and C1q, inducing astrocytes to acquire a pro-inflammatory, and neuron-killing phenotype ^27^. Recognizing the critical role of microglia in responding to environmental challenges and maintaining central homeostasis, we hypothesized that these cells may play a critical role in mediating the detrimental effects of VOC exposure, especially in the context of hypothalamic regulation of metabolism.

To address this gap in knowledge, our current study employs a multifaceted approach, integrating continuous real-time glucose monitoring (CGM), comprehensive hypothalamic and microglial transcriptomics, and the use of novel global and adult-onset microglia-specific mice models of IKK/NF-κB pathway loss of function. Our study establishes, for the first time, microglial IKK/NF-κB signaling as a key mediator of the adverse metabolic effects triggered by VOC exposure.

## Results

### Associations between environmental benzene exposure and metabolic diseases in human; a meta-analysis

To explore the connections between environmental benzene exposure and metabolic health outcomes, we conducted a rigorous meta-analysis of clinical studies. This analysis specifically focused on studies that measured urinary benzene metabolites (particularly trans, trans-muconic acid: t,t-MA) and provided adjusted odds ratios (OR) concerning the risk of metabolic disease. Our comprehensive literature search employed the PubMed research database and included key terms such as (“volatile organic compound” OR “benzene exposure”) AND (“metabolic disease” OR “insulin resistance”) in the abstract or title. This search yielded a total of 128 articles. We excluded 68 articles due to the absence of relevant metabolic measurements, 38 articles for being methodological in nature, and 14 articles for utilizing animal models. Eight studies met our stringent criteria including measurements of urinary t,t-MA levels with readjusted odds ratios (OR) for metabolic disease risk, using parameters such as blood glucose, insulin, or cholesterol levels. The chosen studies encompassed a total of 26,313 subjects, spanning various demographics including pregnant women, children, young adults, adults and elderly individuals ^6–8,10,28–31^. As indicated in the forest plot (Figure 1), we found significant effects of environmental benzene exposure, at levels pertinent to highly polluted urban settings, on the risk of developing metabolic disease. Notably, higher levels of benzene exposure were directly associated with an increased risk of insulin resistance, with a calculated Meta-OR of 1.47, 95%CI [1.33, 1.63], p< 0.001. These data clearly justify the following preclinical study, which was aimed at understanding the molecular mechanisms underlying benzene-induced metabolic disorders.

**Figure 1.**
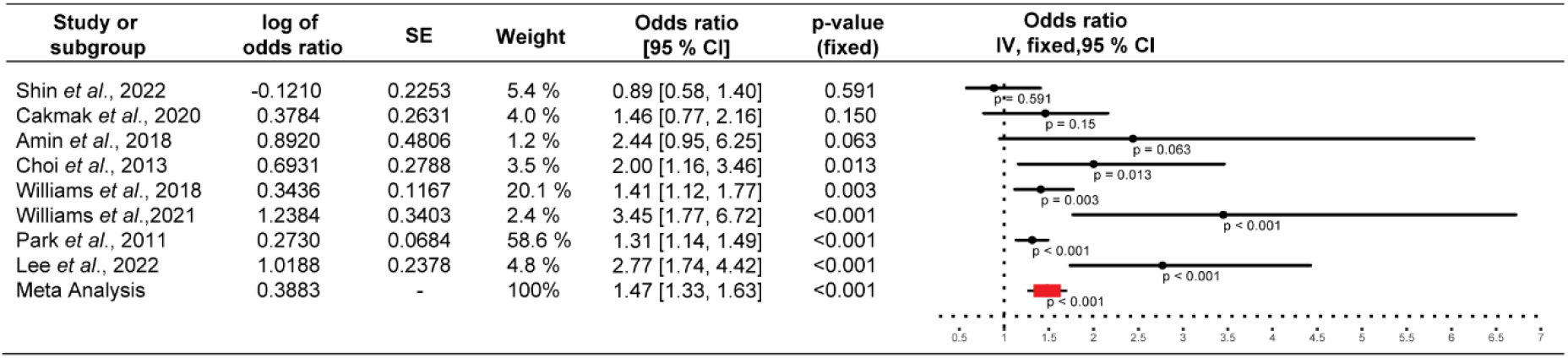
Benzene exposure in human is associated with metabolic disease. Forest plot of the association between benzene exposure and insulin resistance (HOMA-IR). The p-value for the meta-analysis is calculated based on the fixed-effect model.

### Benzene exposure induces rapid hyperglycemia and impairs energy homeostasis in mice

In a previous study, we demonstrated that a 4-week exposure to benzene at levels mimicking smoking (50 ppm) triggered hyperglycemia and hyperinsulinemia, specifically in male mice. However, the temporal impact of this exposure on metabolic outcomes in male mice remained unclear. In the current study, to follow early metabolic changes, we employed a continuous glucose monitoring (CGM) system, enabling real-time tracking of blood glucose levels. After CGM implantation, male mice underwent a 3-day acclimatization period (Figure 2A). We continuously measured blood glucose levels over a course of 7 days. No significant changes in blood glucose levels were detected at the initial 3 days of exposure (Figure S1A-C). However, starting from day 4 of exposure, a significant elevation in blood glucose levels was observed in mice exposed to benzene (p<0.001) (Figure 2B and S1D-F). After 7 days of exposure, a substantial increase of about 20% change (p<0.01) of blood glucose levels compared to day 1 was observed (Figure 2C). The hypothalamus integrates peripheral metabolic inputs and is primarily responsible for regulating energy metabolism. Accordingly, we observed significant reductions in energy metabolism parameters (VO_2_ consumption, VCO_2_ production, respiratory exchange ratio (RER), and heat production) specifically during the dark cycle, after the initial 6 hours of benzene exposure (Figure 2D-G), indicating the involvement of hypothalamic regulation following benzene exposure. Notably, no differences in food intake or total activity levels were detected (Figure 2H and I). The alterations in energy homeostasis parameters remained evident even 7 days after the 4-week exposure in male mice (Figure S1G and H) indicating the prolonged effects of benzene exposure on regulation of energy metabolism. Intriguingly, benzene exposure led to increased serum corticosterone levels (measured either after 24 hours or on day 4 of exposure) in male mice (Figure S1I), suggesting a link between short-term stress response mechanisms and elevated blood glucose levels. However, the levels of corticosterone were normalized after 4-week exposure indicating the involvement of other non-stress related mechanisms to the disruptions in energy homeostasis (Figure S1I). Notably, no differences in energy homeostasis (Figure S2A-D), body weight or glucose tolerance were observed between females exposed to benzene and the control group ^18^.

**Figure 2.**
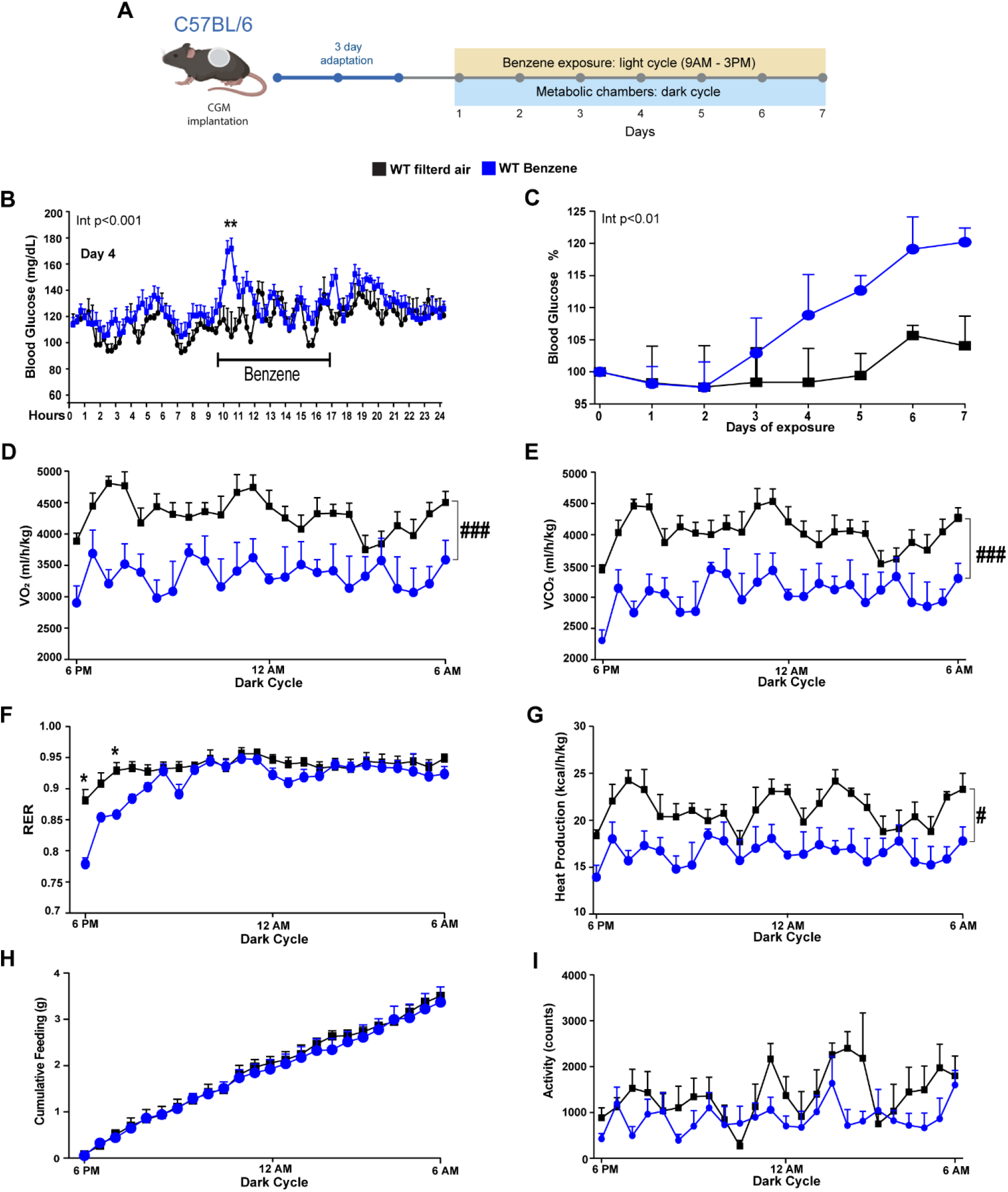
Rapid metabolic imbalance induced by benzene exposure in mice. (A) Experimental design illustrating the monitoring of glycemic changes and energy homeostasis in 12-week-old WT (C57BL/6J) male mice exposed to benzene (50 ppm). A transdermal sensor for continuous glucose monitoring (CGM) was implanted, and following a 3-day acclimatization period, animals were exposed for 6 hours during the light cycle, from 9 AM to 3 PM, over a 7-day period; (B) Day 4 of CGM; (C) Percentage change in blood glucose levels, calculated from day 0 (pre-exposure) to day 7. Energy homeostasis parameters measured during the dark cycle after 1 day of exposure (D) Oxygen consumption (VO2) consumption [ml/h/kg]; (E) Carbon dioxide production (VCO2) [ml/h/kg]; (F) Respiratory exchange ratio (RER); (G) Heat production [kcal/h/;kg]; (H) Cumulative food intake (g); (I) Locomotor activity. Error bars represent the standard error of the mean (SEM) for n = 8-10 mice/group. All values are presented as mean ± SEM. Repeated measure ANOVA (^#^p<0.05, ^###^p<0.001) and the Newman-Keuls post hoc test (*p< 0.05, **p<0.01). The interaction effect of repeated measure ANOVA (exposure versus time) is indicated on the graph.

### Benzene exposure impairs hypothalamic insulin responsiveness through transcriptional alterations

The hypothalamus plays a crucial role in the regulation of energy expenditure and glucose levels, which can be influenced by a variety of environmental factors ^32^. To assess the impact of benzene-induced metabolic shift on hypothalamic transcriptome, we conducted an analysis of differential gene expression in the hypothalamus of male mice acutely exposed to benzene for 24 hours, comparing them to filtered-air exposed mice using bulk RNA-sequencing (RNAseq). The RNA-seq analysis revealed 737 differentially expressed genes in the whole hypothalamus of benzene-treated mice. Among these, 405 transcripts exhibited significant upregulation, while 332 were significantly downregulated when compared to controls (adjusted p <0.1, log2 fold-change>1) (Figure 3A). Remarkably, one of the most enriched pathways in response to benzene exposure was associated with the response to insulin, as indicated by gene ontology (GO) pathways and gene set enrichment analysis (GSEA) (Figure 3B).

**Figure 3.**
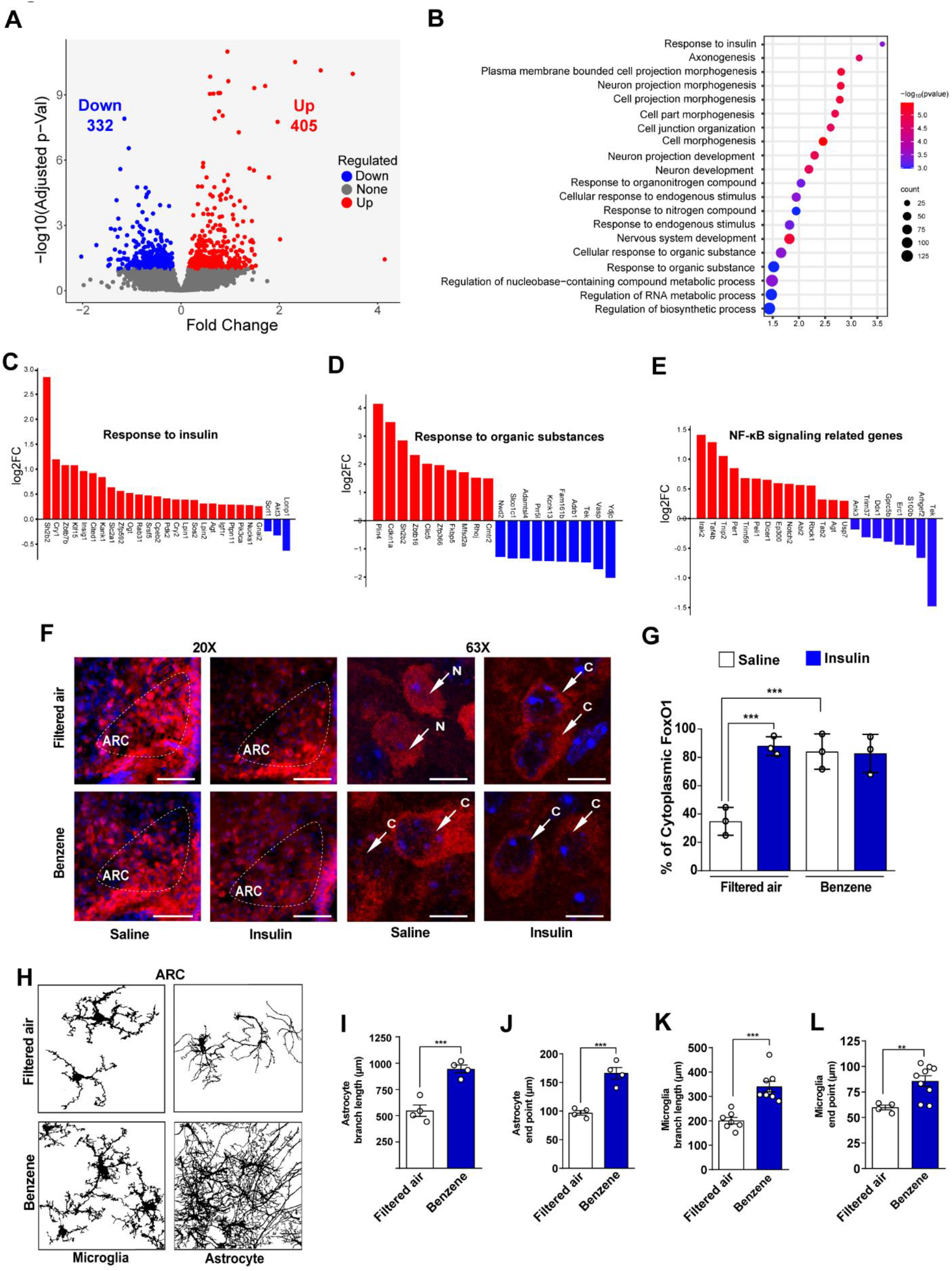
Rapid modification of hypothalamic transcriptome and impairment of hypothalamic insulin signaling and glial morphology induced by benzene exposure. Analysis of the hypothalamic transcriptome following acute benzene exposure. (A) Volcano plots depict differentially expressed genes (FDR < 0.1), with upregulated genes in red and downregulated genes in blue, comparing benzene-exposed and filtered air-exposed hypothalamus; (B) Gene ontology analysis shows enriched pathways in the hypothalamus after benzene exposure; (C) Upregulated and downregulated genes in response to insulin; (D) Upregulated and downregulated genes in response to organic substances; (E) Upregulated and downregulated genes in NF-kB signaling, determined using DESeq2 analysis, in 12-week-old male mice exposed to benzene versus control, with adjusted p < 0.1 and log2 fold-change > 0.1; (F) Representative immunostaining images of FoxO1 (red) and DAPI (blue) in the arcuate nucleus of the hypothalamus (ARC) in 12-week-old male mice exposed to benzene for 6 hours, euthanized 15 minutes after insulin injection (i.p.; 3 I.U./kg B.W.) or saline. White arrows in the confocal images (63x) indicate FoxO1 protein localization (C indicates cytoplasmic, N indicates nuclear). Scale bars: 200 μm and 10 μm in the merged picture; (G) Percentage of cytoplasmic FoxO1 expression in the ARC, with error bars representing SEM for n = 3-4 mice per group. Images were analyzed from at least three to four sections; (H) Skeleton images representing astrocyte and microglia morphology in the ARC of male mice exposed to benzene or filtered air; (I) Quantification of astrocyte branch length (µm) and (J) astrocyte endpoints (µm). (K) Quantification of microglia branch length (µm) and (L) microglia endpoints (µm), with error bars indicating SEM for n = 4-8 mice per group. Two-way ANOVA with the Newman-Keuls post hoc test (**p < 0.01, ***p < 0.001).

Among the key genes that were upregulated in response to benzene exposure and are involved in insulin regulation were *Sh2b2, Cry1, Cry2, Zbtb7b,Klf15, Cited1, Slc2a1, Ogt, Rab31, Srsf5, Cpeb2, Pdk2, Lpin1, Sos2, Agt, Ptpn11, Pik3ca* and *Nucks1* (Figure 3C).Some of these genes have been demonstrated to stimulate insulin signaling, while others are involved in insulin-induced activation of mitogen-activated protein (MAP) kinases mediated by Ras ^33^. Conversely, *Akt3, Lonp1* and *Sorl1* were found to be downregulated (Figure 3C). In line with the upregulation of cellular responses to organic substances (Figure 3D), we observed an upregulation of certain genes related to the inflammatory response within the NF-kB signaling pathway, including *Irak2, Taf4b, Tnip2, Per1, Trim59, Peli1, Dicer1, Agt and Usp7*. Conversely, inflammatory genes such as *Tek, S100b, Gprc5b* and *Trim37* were downregulated in the hypothalamus of mice exposed to benzene (Figure 3E).

To investigate the potential impact of benzene exposure on hypothalamic insulin signaling, male mice were acutely exposed to benzene for 6 hours while fasting, and brains collected after receiving a single dose of insulin (i.p.; 3 I.U./kg B.W.) or saline injection. In control mice, insulin-stimulated FoxO1 phosphorylation led to an increase in its cytoplasmic translocation in both the arcuate nucleus (ARC) (Figure 3F and G) and the ventromedial nucleus of the hypothalamus (VMH) (Figure S3A-C). In contrast, following benzene exposure, there was a significant increase in cytoplasmic FoxO1 levels (p<0.001), which did not exhibit further responsiveness to insulin (Figure 3G and Figure S3A-C). A similar response pattern to benzene exposure was also observed with MAPK signaling, a pathway implicated in the development of insulin resistance ^34^. In control mice, insulin stimulation increased MAPK phosphorylation (Figure S3D-F). However, in mice exposed to benzene, p-MAPK was significantly elevated as compared to unstimulated control (p<0.001), and did not further respond to subsequent insulin stimulation (Figure S3D-F). Consistent with our prior findings ^18^, we found significant alterations in the morphology of astrocytes and microglia within the hypothalamus of mice exposed to benzene. Specifically, we observed a notable increase in the extension of branches and endpoints of these cells (p<0.001) (Figure 3H-L) suggesting a higher level of activity among microglia and astrocytes, possibly signifying cellular hypertrophy and an adaptive immune response of these glial cells to the presence of benzene.

### Benzene-induced metabolic imbalance is caused by activation of IKKβ/NF-κB signaling in immune cells

Disrupted insulin signaling and increased inflammation mutually contribute to the progression of metabolic disease ^12,13,14^. We have previously demonstrated that prolonged benzene exposure induced systemic inflammatory responses in male but not female mice ^18^. To test the possible causality between benzene-induced central inflammation and impaired systemic glucose metabolism, we ablated the prototypical proinflammatory IKKβ/NF-κB signaling pathway in immune cells. We used the fractalkine receptor (Cx3cr1)-driven Cre, which is widely expressed in immune cells in the body including microglia, on ROSA26-GFP background and crossed them with IKKβ^lox/lox^ mice ^35–37^. In control (IKK^flox/flox^) mice acute, 6-hour exposure to benzene significantly increased TNFα expression in microglia, with approximately 80% of GFP^+^ stained hypothalamic microglia showing TNFα production (GFP^+^/TNFα^+^), indicating their inflammatory status ^36^. In contrast, Cx3cr1^GFPΔIKK^ male mice exposed to benzene had similar numbers of GFP^+^/TNFα^+^ microglia as filtered air control mice (Figure 4A-B), and no aberrations in microglia morphology as compared to in benzene-exposed controls (Figure 4D-F), suggesting the absence of microglial activation. As previously, the numbers of microglia were not increased by acute exposure in control animals ^18^, but similarly reduced in unexposed or exposed Cx3cr1^GFPΔIKK^ mice compared to control (Figure 4C). Accordingly, while benzene-exposed control mice exhibited increase in GFAP^+^ astrocytes numbers and alterations in astrocytes’ morphology characterized by the numbers and length of GFAP positive processes within the hypothalamus, astrocyte in benzene-exposed Cx3cr1^GFPΔIKK^ male mice was not significantly affected (Figure. S4). Finally, Cx3cr1^GFPΔIKK^ mice exposed to benzene during the light cycle exhibited similar VO_2_ and VCO_2_ as filtered air-exposed controls compared to benzene-exposed control mice (Figure 4G and H). Additionally, while benzene exposure induced glucose intolerance in control mice, Cx3cr1^GFPΔIKK^ male mice exhibited normal glucose tolerance, conducted after 4 weeks of benzene exposure (Figure 4I). Further, insulin stimulation promoted similar responses in benzene-exposed Cx3cr1^GFPΔIKK^ male mice, indicated by FoxO1 translocation to the cytoplasm, comparable to control Cx3cr1^GFPΔIKK^ mice (Figure S4A-C), with appropriate insulin-stimulated increase in MAPK phosphorylation (Figure S4D-E). Thus, preventing the activation of immune cells in response to benzene exposure protects from benzene-induced metabolic imbalance.

**Figure 4.**
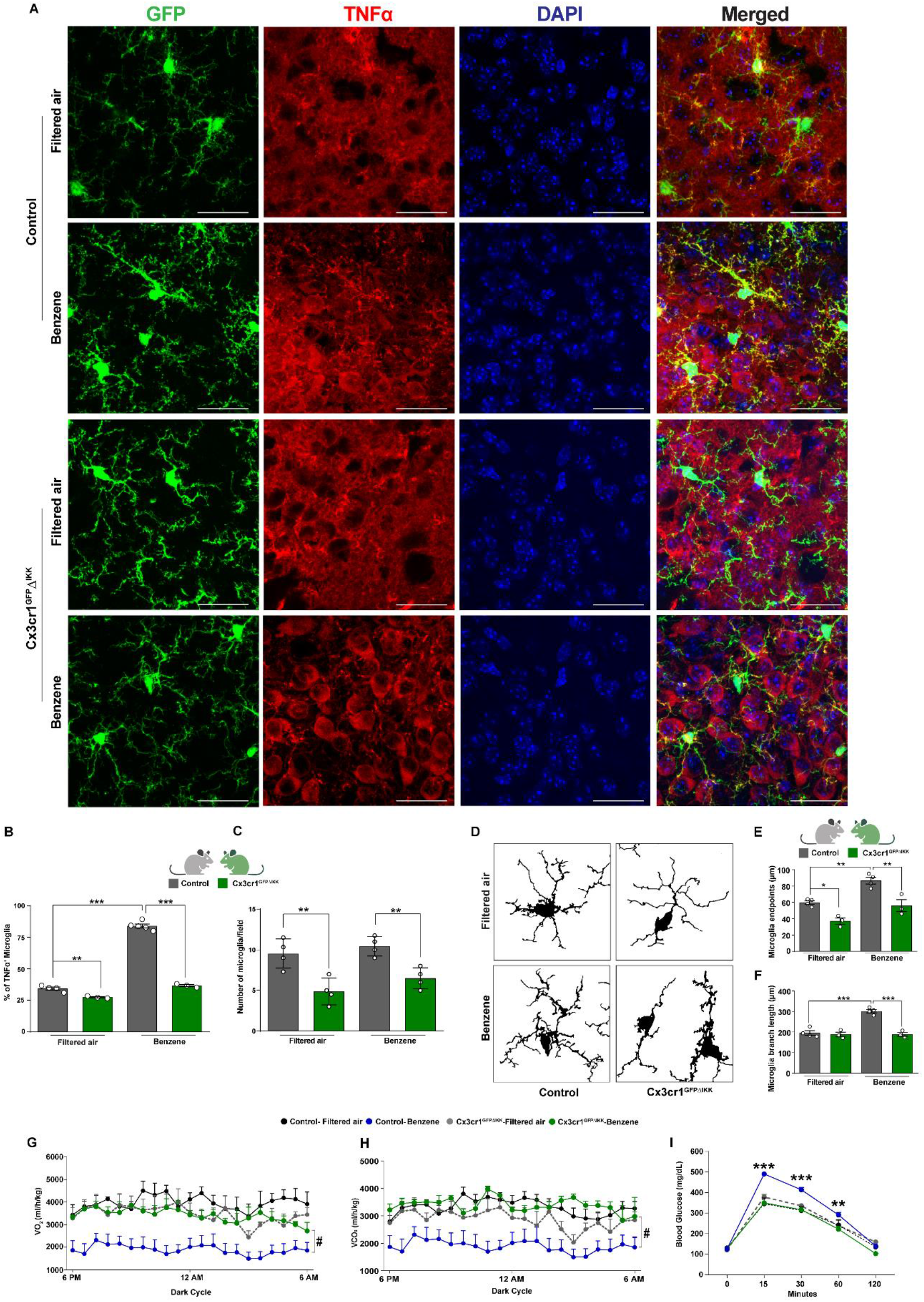
Restoration of benzene-induced metabolic imbalance through IKKβ/NF-κB signaling depletion in immune cells. (A) Representative images demonstrating TNFα staining (in red), DAPI (in blue), and microglia (GFP) in the arcuate nucleus of the hypothalamus (ARC) of 12-week-old male mice (gray, IKK^fl/fl^; green, Cx3cr1^GFPΔIKK^). Scale bars: 10 µm; (B) Percentage of microglia positive for TNFα in the ARC (gray, IKK^fl/fl^; green, Cx3cr1^GFPΔIKK^); (C) Quantification of GFP^+^ microglia; (D) Skeleton images representing microglia morphology in the ARC of Cx3cr1^GFPΔIKK^ mice exposed to benzene compared to filtered air-exposed animals; (E) Measurement of microglia endpoints; (F) Microglia branch length (µm); Impact of 6-hour benzene exposure during the light cycle on energy homeostasis parameters in Cx3cr1^GFPΔIKK^ and control mice measured during the dark cycle (G) Oxygen consumption (VO2) [ml/h/kg]; (H) Carbon dioxide production (VCO2) [ml/h/kg]; (I) Glucose tolerance test (GTT) in Cx3cr1^GFPΔIKK^ mice and their control IKK^fl/fl^ littermates exposed for 4 weeks/6 hours per day. Error bars represent the standard error of the mean (SEM) for n = 4-8 mice per group. Two-way ANOVA with the Newman-Keuls post hoc test (*p < 0.05, **p < 0.01, ***p < 0.001). Repeated measure ANOVA (#p < 0.05).

### Benzene exposure induces transcriptional remodeling in microglia

Activation of the central immune system, specifically microglia, under stress conditions was suggested to play a critical role in driving peripheral hyperglycemia and impaired energy balance ^38–40^. To specifically determine the role of central inflammatory responses to benzene exposure, we used CD11b^+^ magnetic-activated cell sorting (CD11b-MACS) to isolate microglia from the brains of adult male mice exposed to benzene or filtered air for 24 hours and performed bulk RNA-seq analysis (Figure 5A). The purity of the CD11b-MACS positive microglia was verified by flow cytometry (Figure S5A). The CD11b-MACS positive fraction was composed of ~ 80% CD11b^+^CD45^+^ cells, as compared to 10–20% of the input (Figure S5B, two-way ANOVA, **p < 0.01). In addition, because CD45 is expressed by non-microglial cells, including monocytes and macrophages, we further assessed the purity of samples with partial deconvolution method, CIBERSORTx ^41^. CIBERSORTx analysis indicated an average of 2.1% non-microglial (astrocytes, neurons, oligodendrocytes) cells per sample (Figure S5). RNA-seq analysis showed 1531 differentially expressed genes in microglia of benzene-treated mice, where 749 transcripts were significantly elevated and 782 were significantly reduced as compared to control (adjusted p < 0.1, |log2 fold-change|>1) (Figure 5B). Based on our analysis of gene ontology (GO) pathways by gene set enrichment analysis (GSEA), exposure to benzene elicited a significant upregulation of pathways related to chemotaxis, response to nicotine, generation of metabolites and energy, cellular metabolic processes, energy derivation by oxidation of organic compounds, and cytokines production in microglia. Notably, these upregulated pathways were driven by genes associated with cellular responses to organic chemicals, immune response, and the regulation of NF-kB transcription factor activity (Figure 5C). Among the key genes with significant upregulation in response to benzene exposure were *Myd88, Nod2, Trim12c, Ltf, App, Amfr, Ikbkg, Btrc, Npm1* and *Nkiras1* (Figure 5D). These genes have been shown to activate NF-κB signaling ^42–46^ and have been linked to proinflammatory stimuli and neurodegenerative diseases ^47^. Conversely, certain genes related to the inflammatory response within the NF-kB signaling, such as *Klf4, Il1b, Trim39 and Nfkbib,* were downregulated in microglia from benzene-exposed mice. Similarly, we observed upregulation in the expression of chemotaxis genes such as *Retnlg, Mmp28,* and *Bsg*, which are known to influence the polarization of microglia toward an anti-inflammatory state ^48–50^ (Figure 5E). On the other hand, genes like *Ndufa13, Anxa1, H13,* and *Nucb2*, associated with the regulation of inflammation activation and cytokine responses, also displayed upregulation ^51–54^ (Figure 5F). These findings collectively suggest an influence of several concurrent homeostatic pathways in response to benzene exposure and provide further evidence supporting the notion that acute benzene exposure prompts a shift in microglia toward a reactive phenotypic state, with the central involvement of the NF-κB signaling pathway.

**Figure 5.**
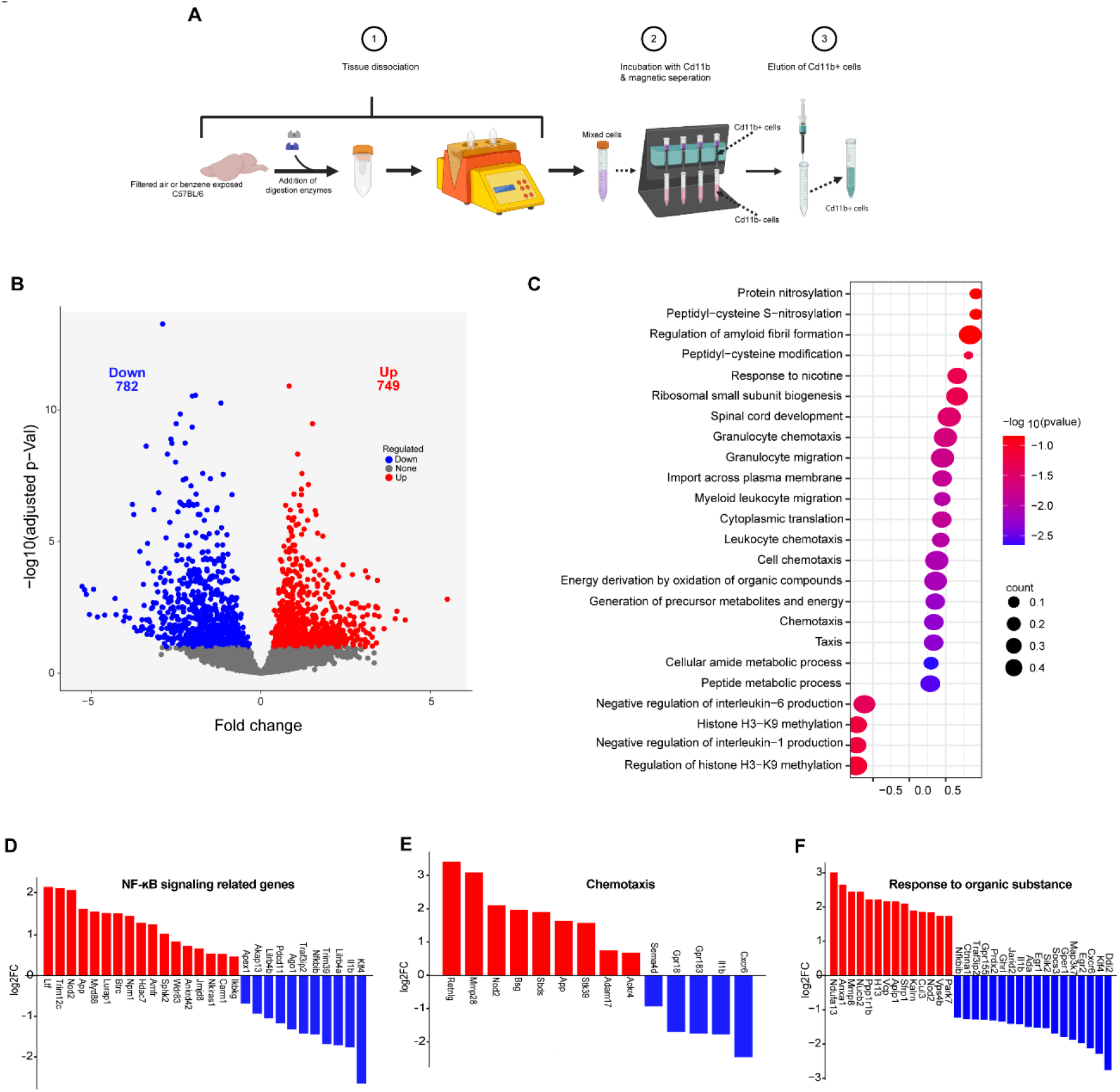
Acute benzene exposure induces microglial transcriptome remodeling toward an inflammatory activated state. (A) Schematic showing experimental strategy for the isolation and purification of microglia from control and 24-h benzene-exposed male mice; (B) Volcano plots illustrating differentially expressed genes (FDR < 0.1), with upregulated genes in red and downregulated genes in blue, when comparing microglia from benzene-exposed and filtered air-exposed mice; (C) Gene Ontology analysis shows enriched pathways in the microglia after benzene exposure; (D) Genes upregulated (in red) and downregulated (in blue) in the NF-kB signaling pathway; (E) Genes differentially upregulated (in red) and downregulated (in blue) in chemotaxis; (G) Genes differentially upregulated (in red) and downregulated (in blue) in response to organic substances, determined using DESeq2 analysis in 12-week-old male mice exposed to benzene versus control (n=5-6 mice per group).

### Activation of microglial IKKβ/NF-κB signaling induces metabolic imbalance upon benzene exposure

With the rapid activation of NF-κB signaling-related genes in the microglia transcriptome following exposure to benzene, we hypothesized that NF-κB pathway activation specifically within microglia could be critical for the observed hyperglycemia and insulin resistance caused by benzene ^18^. To test this hypothesis, we took advantage of the recently developed tamoxifen-inducible Tmem119-Cre^ERT^^2^ mice on ROSA26-EGFP background ^55^ and crossed them with IKKβ^lox/lox^ mice. 14-weeks-old male mice lacking IKKβ exclusively in microglia (TMEM119^ERΔIKK^) displayed normal body weight and blood glucose levels, comparable to those of their control littermates (Figure S6A-B). Remarkably, the TMEM119^ERΔIKK^ males were completely resistant to benzene-induced microgliosis, as evidenced by the minimal TNFα production observed in the hypothalamic microglia of TMEM119^ERΔIKK^ mice (Figure 6A and B). Furthermore, there were no detectable changes in energy homeostasis parameters (VO_2_ and VCO_2_,) in TMEM119^ERΔIKK^ mice, compared to control animals, whether exposed to air or benzene (Figure 6C and D). Remarkably, the TMEM119^ERΔIKK^ male mice remained protected against benzene-induced hyperglycemia, after 4-weeks of benzene exposure while maintaining euglycemia (Figure 6E). Furthermore, the elevated expression of inflammatory genes related to NF-kB signaling and insulin responsive genes detected in the hypothalamic transcriptome of benzene-exposed control mice was mostly restored in TMEM119^ERΔIKK^ mice, reflecting air-exposed controls (Figure 6F and G). Thus, the microglia-specific ablation of NF-κB pathway provides protection against benzene-induced metabolic disturbances, highlighting a potential for therapeutic interventions.

**Figure 6.**
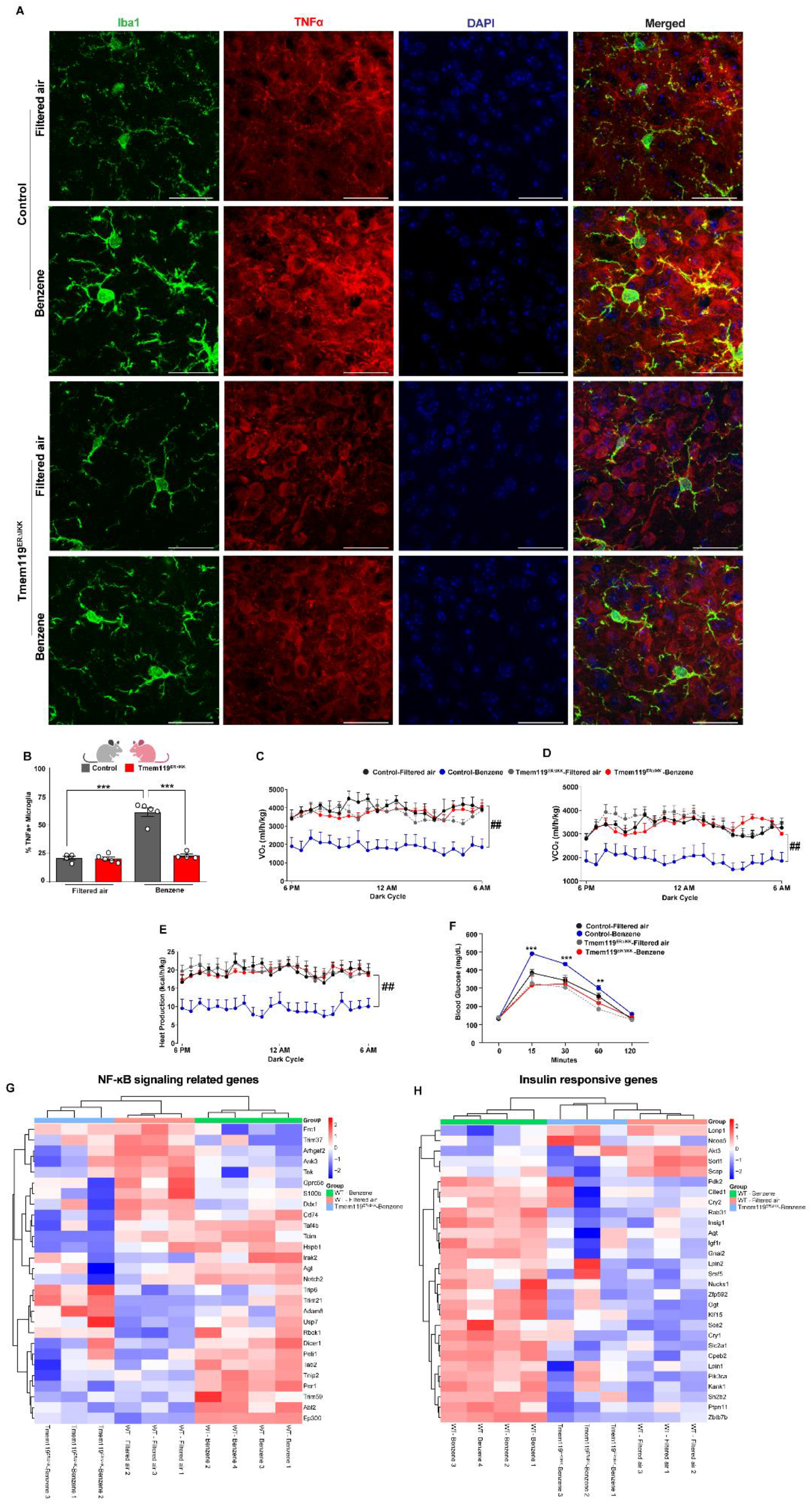
Deletion of microglial IKK/NF-κB signaling protects against benzene-induced metabolic imbalance. (A) Representative images demonstrating staining for TNFα (in red), microglia (Iba-1, in green), and DAPI (in blue) in the arcuate nucleus of the hypothalamus (ARC) of tamoxifen-inducible TMEM119-IKK knockout (TMEM119^ERΔIKK^) male mice. Scale bars: 10 µm; (B) Percentage of TNFα-positive microglia in the ARC (gray, IKK^fl/fl^; red, TMEM119^ERΔIKK^). Energy homeostasis parameters measured during the dark cycle in TMEM119^ERΔIKK^ and control mice following 6 hours of benzene exposure: (C) Oxygen consumption (VO_2_) [ml/h/kg]; (D) Carbon dioxide production (VCO_2_) [ml/h/kg]; (E) Heat production [kcal/h/;kg]; (F) Glucose tolerance test (GTT) in TMEM119^ERΔIKK^ mice and their control IKK^fl/fl^ littermates exposed for 4 weeks/6 hours per day, compared to control and TMEM119^ERΔIKK^ filtered air-exposed 12-week-old male mice. Error bars represent the standard error of the mean (SEM) for n = 4-8 mice per group. Two-way ANOVA with the Newman-Keuls post hoc test (**p < 0.01, ***p < 0.001). Repeated measure ANOVA (##p < 0.01). Analysis of the hypothalamic transcriptome following acute benzene exposure of TMEM119^ERΔIKK^ and control male mice as compared to filtered air-exposed 12-weeks old control male mice. Heat maps of top genes related to (F) response to insulin, and (H) NF-kB signaling in the hypothalamus. n=3/4 mice per group.

## Discussion

Utilizing a continuous glucose monitoring (CGM) system enabled us to demonstrate that a short-term exposure to benzene of 6 hours/day for four days, leads to elevated blood glucose levels and rapid disruptions in energy homeostasis in male mice. Importantly, we found that hypothalamic insulin responsiveness and activation of microglia play a significant role in these metabolic disturbances following benzene exposure. Further, we uncovered the underlying molecular mechanism showing that NF-κB pathway in microglia plays a central role in the initial neuroinflammatory response to benzene exposure. These data indicate that inhibiting microglial immune response to environmental stressors, such as VOCs, could offer potential therapeutic benefits in treating the consequent metabolic impairments.

Our comprehensive meta-analysis, which covers a wide range of demographics such as the elderly, children, young adults, adults, and pregnant women ^6–8,10,28–31^, shows a strong positive relationship between levels of trans, trans-muconic acid (t,t-MA), benzene urinary metabolites, and indicators of insulin resistance and blood glucose levels. These data are in line with recent findings from a comprehensive cross-sectional study utilizing data from The National Health and Nutrition Examination Survey showed a strong correlation between VOCs exposure and development of insulin resistance within the broader population ^56^. Together, our meta-analysis along with the aforementioned studies suggest that exposure to benzene, at levels relevant to the general population, is a potential risk factor for developing diabetes and insulin resistance.

We addressed the effects of benzene on peripheral metabolism and neuroinflammation after short- and long-term exposure. Our previous study had shown that a chronic 4 week-exposure to benzene at levels resembling smoking, induced insulin resistance, hyperglycemia, and hypothalamic inflammation ^18^. Here we show that benzene exposure for only 6 hours during the light cycle, led to immediate decrease in parameters of energy homeostasis that were evidenced during the following dark cycle. Elevated blood glucose levels were detected by CGM on the 4th day of exposure. These changes were accompanied by an increase in glucocorticoid levels in the blood, suggesting the rapid involvement of the hypothalamic-pituitary-adrenal (HPA) axis ^57^. Interestingly, the reductions in parameters of energy expenditure were evident even a week after the cessation of 4 week-exposure, indicating the prolonged effects of benzene exposure on energy metabolism. However, glucocorticoid levels were normalized by that time, indicating the impact on specific hypothalamic circuits that involve multiple signaling processes on hypothalamic plasticity ^58^.

The rapid changes in energy and glucose metabolism correlated with hypothalamic RNA-seq analysis conducted 24 hours after exposure to benzene. Notably, the most enriched GO terms in the hypothalamic transcriptome of benzene-exposed mice, compared to controls, were related to responses to insulin. Particularly interesting was the significant upregulation of Src homology 2B (SH2B) family member, *Sh2b2*, which exhibited an almost threefold increase in expression. The role of SH2B2 in insulin action is multifaceted. SH2B2 overexpression has been shown to prolong insulin-stimulated tyrosine phosphorylation of insulin receptors and IRS proteins ^59^. Conversely, SH2B2 can promote the ubiquitination and internalization of insulin receptors, inhibiting insulin signaling ^60,61^. It may also negatively regulate insulin signaling through its interaction with SOCS family members ^62^. Additionally, multiple cytokines can activate SH2B2 via JAK family members ^63^. Notably, the deletion of SH2B2 has been found to increase insulin sensitivity in mice ^64^. In line with this, our analysis detected the upregulation of hypothalamic genes such as *Cited1, Cry1, Cry2, Agt, Rab31, Sos2,* and *Ptpn11*, which have previously been implicated in the activation of the RAS–MAPK pathway and insulin resistance ^65–69^. RNAseq was further supported by immunohistochemistry showing that benzene exposure rapidly impaired basal and insulin-stimulated FoxO1 cellular translocation in the hypothalamus and led to elevations in hypothalamic MAPK signaling, indicating impaired hypothalamic insulin signaling ^70^. Interestingly, similar hypothalamic transcriptional changes involved in insulin action, and inflammation were detected with exposure to particulate matter smaller than 2.5 μm in size (PM_2.5_) over a 14-week period ^71^. Identifying the molecular mechanisms underlying these rapid changes will require further mechanistic investigation of the roles of specific molecules and other signaling pathways in the hypothalamus.

VOCs can reach the CNS in just a few minutes through direct transport or via systemic inflammation initiated in the lung tissue immune cells ^72,73^. However, the molecular mechanism by which VOCs trigger neuroinflammation remains insufficiently understood. *In vitro* studies using rat models or human cell lines have shown that exposure to VOCs triggers oxidative stress response accompanied by the generation of inflammatory mediators ^74^. Oxidative stress was shown to be a key player in activating NF-kB through the phosphorylation of its associated IkB proteins, underlying the principal mechanism for cigarette smoke-induced tissue damage ^75^. We have previously demonstrated elevated expression levels of *Ikbkb*, *Ikbke, Tnf*, *Il1b*, and *Il6* genes in the hypothalamus and the liver of male mice exposed to benzene for 4 weeks ^18^. This was associated with increased gene expression of the benzene metabolite Cyp2e1, confirming benzene-induced activation of CYP2E1 in glial cells ^76^. The generation of Cx3cr1^GFPΔIKK^ mice, in which IKK was eliminated from immune cells, including monocytes, macrophages, and microglia, yielded further compelling evidence. These mice showed protection against alterations in glucose metabolism and energy expenditure induced by benzene exposure. This observation strongly implies that the NF-kB signaling in immune cells plays a pivotal role in modulating peripheral metabolism. Within the CNS, the activation of IKKβ/NF-κB in hypothalamic microglia and astrocytes has been linked to metabolic imbalance in obese mice ^39,38,77^. Our research reveals that microglia are particularly susceptible to environmental factors such as benzene. It offers direct evidence linking the IKKβ/NF-κB signaling pathway in microglia to peripheral insulin resistance and disrupted energy balance.

We observed significant alterations in the microglial transcriptional profile following 24 hours of benzene exposure, with a distinct shift towards an inflammatory phenotype upon exposure to benzene. This was evidenced by elevated expression of genes associated with chemotaxis, cellular response to chemicals, and notably, NF-κB signaling. Interestingly, some genes significantly influenced multiple enriched pathways. Noteworthy among these genes, the *APP* gene, known for its regulation of microglial proinflammatory responses in Alzheimer’s disease (AD) ^78^, and the *Nod2* gene, which amplifies microglial inflammation via the TAK1-NF-κB pathway ^79^, displayed significant upregulation. Conversely, negative regulators of *IL-1β* and *IL-6* production were downregulated. Additionally, the downregulation of *Klf4*, a regulator of microglial activation and neuroinflammation ^80^, and NF-κB inhibitor beta (*Nfkbib*) gene were observed in benzene-exposed microglia. Although KLF4 is associated with neuroinflammation through the regulation of IL-1β, cyclooxygenase-2 (Cox-2), and inducible nitric oxide synthase (iNOS) production ^80^, its expression is reduced in cancer ^81^. NFKB1, a pivotal driver of cellular inflammation and immunity, encodes the p105 subunit of NF-κB, which is processed to generate the NF-κB p50 subunit ^82^. Simultaneously, genes related to NF-κB signaling activation, such as *Trim12c*, *Ltf*, *Npm1, Btrc* and *Nkiras1* ^42–46^ exhibited upregulation. Furthermore, the upregulation of the *MyD88* gene, essential for TLR4/MyD88/NF-κB signaling and the production of TNF-a and IL-6 ^42^, and the *Amfr* gene, a regulator of NF-κB signaling within the cGAS–STING pathway ^83^, suggest the possible involvement of multiple upstream regulators acting synergistically to drive microglial activation in response to acute benzene exposure.

The distinct microglial inflammatory response to benzene exposure prompted us to generate the microglia specific IKK-knockout mice (TMEM119^GFPΔIKK^). These mice showed a blunted neuroinflammatory response and no metabolic impairments such as insulin resistance or imbalance in energy expenditure following benzene exposure. Furthermore, we observed a limited overlap in hypothalamic genes regulated by responses to insulin or NF-kB signaling between the control group and TMEM119^GFPΔIKK^ animals exposed to benzene. For example, hypothalamic genes upregulated by benzene exposure, such as *Sh2b2, Cited1*, *Cry1*, *Cry2*, *Rab31*, *Sos2* and *Ptpn11*, exhibited normalization in the hypothalamus of benzene-exposed TMEM119^GFPΔIKK^ mice, bringing their expression levels in line with those observed in non-exposed control mice. These data, using the TMEM119^GFPΔIKK^ mice, provide direct evidence for the involvement of NF-κB-induced microglial activation in the impairment of peripheral metabolism.

Our data are in line with studies showing that inhibiting microglial inflammation can protect against obesity and reduce microgliosis in the hypothalamus ^84,38^, and that selective depletion of IKKβ in microglial cells results in reduced brain expression of the NF-κB target genes, IL-1β, IL-6 ^85^. A recent paper, however, suggested that the neuroinflammatory response in microglia provides a protective mechanism against insulin resistance. Using the *Cx3cr1*^CreER/+^::*Ikk*^fl/fl^ inducible mouse model fed high-fat diet (HFD), the authors showed that loss of microglial IKKβ protected the mice from weight gain, however paradoxically it worsened glucose tolerance. Conversely, stimulating microglial inflammatory signaling improved glucose tolerance ^86^. More research will be required to understand the discrepancy between those findings and the current study. It is possible however, that microglial response to HFD feeding differs from their response to environmental stressors. That is especially intriguing since the effect of microglial signaling on glucose homeostasis was body weight-independent ^86^. On the other hand, while we did not observe any differences in body weight over the period of 4-weeks of benzene exposure ^18^, TMEM119^GFPΔIKK^ mice were protected from benzene-induced hyperglycemia. CNS glucose responsiveness is governed by an extensively distributed network of glucose-sensing neurons spanning various brain regions ^19^. This strongly suggests that microglial regulation of glucose tolerance in response to environmental stressors engages multiple neuronal subtypes within the brain. Further investigations are essential to elucidate these variations in microglial responses.

### Limitations of the study

Our study has a few limitations including the use of male mice only that was justified by our previous study detecting significant sex specific effects of benzene exposure. Given that metabolic changes were seen only in male mice, we restricted our analysis to males. However, it is worth noting that there may exist more subtle metabolic responses in both males and females that differ from the sexually dimorphic outcomes related to glucose tolerance, insulin responsiveness, and energy homeostasis. An additional comparison between male and female mice will be needed to fully elucidate sex-specific mechanisms of exposure effects in the CNS. Second, Tmem119-based model is exclusive to the microglia throughout the CNS ^87^, leaving uncertain which microglial region is involved in the response to benzene exposure or what is the role of other glia cell types in the observed phenotype. Third, only one concentration of benzene that mimics cigarette smoking was used ^88^. Additional exposures and mixtures resembling urban environment throughout the lifespan will be useful in understanding how environmental VOC mixtures affect glucose metabolism. Finally, our study identified the role of inflammatory NF-κB signaling in benzene exposure however, it is still possible that other signaling pathways in the CNS or periphery are involved.

In summary, our study provides novel insights into the impact of benzene exposure on neuroinflammation and metabolic imbalance, highlighting the potential contribution of environmental contaminants to the rising prevalence of chronic metabolic health conditions.

## Abbreviations

CNS: central nervous system
CGM: continuous glucose monitoring
NF-κB: nuclear factor kappa B
VOCs: volatile organic compounds
MBH: mediobasal hypothalamus
ARC: arcuate nucleus of the hypothalamus
VMH: ventromedial hypothalamic nucleus
Cx3cr1: C-X3-C motif chemokine receptor 1
t,t-MA: trans, trans-muconic acid

## Acknowledgements

This study was supported by American Diabetes Association 1-lB-IDF-063, CURES Center Grant P30ES020957, NIEHS R01ES033171, NIA RF1AG078170 and CLEAR P42ES030991 for M.S. L.K was further supported by 5T32GM142519-02 and L.S was supported by 5T32HL120822-09. Services and support were provided by Genomic Sciences Core of the Oklahoma Nathan Shock Center, P30AG050911, the WSU Microscopy, Imaging and Cytometry Resources Core P30CA22453 and R50CA251068-01 and WSU Genomics Core.

## Author contributions

L.K.D., H.S.M.J., L.S., A.T.D.S., L.K., R.S and S.S. carried out the research and reviewed the manuscript. U.K. designed the CGM study. A.M performed and analyzed the meta-analysis. M.S. designed the study and analyzed the data. M.S. wrote the manuscript and is responsible for the integrity of this work. All authors approved the final version of the manuscript.

## Declaration of interests

The authors declare no competing interests.

## Resource Availability

Further information and requests for resources and reagents should be directed to and will be fulfilled by the Lead Contact, Marianna Sadagurski. Any additional information required to reanalyze the data reported in this paper is available from the lead contact upon request.

No unique reagents were generated in this study.

## STAR Methods

### Animals

All mice were on C57BL/6J background and maintained under standard laboratory conditions with standard chow low-fat diet (Research diets). All mice were provided with water *ad libitum* and housed in temperature-controlled rooms (22⁰C) on a 12-hour/12-hour light-dark cycle. If animals exhibited any indication of illness or distress, the laboratory staff conferred with on-site veterinary staff immediately to recommend appropriate interventions. Anesthesia for euthanasia was by avertin. The health status checks were conducted regularly in the animal breeding facility. All animal experiments were performed in accordance with NIH guidelines for Animal Care and Use, approved and overseen by Wayne State University Institutional Animal Care and Use Committee (IACUC).

### Rodent lines and experimental animals

C57Bl/6J: Mouse line was purchased from The Jackson Laboratory (strain #000664).

Cx3cr1^Cre^ (GFP): Mouse line was purchased from The Jackson Laboratory (strain #005582).

Tmem119^CreER^: Mouse line was purchased from The Jackson Laboratory (strain #031820).

Ikbkb^fl/fl^: IKKb flox mice were obtained from the laboratory of Dr. Michael Karin ^89^.

Cx3cr1^GFPΔIKK^: Heterozygous Cx3cr1^Cre^ (GFP) mice were bred to be homozygous Ikbkb^fl/fl^. Experimental animals were generated by crossing Cx3cr1^Cre/+^ (GFP)::Ikbkb^fl/fl^ mice with Ikbkb^fl/fl^ mice. Ikbkb^fl/fl^ littermates were used as controls.

Tmem119^ERΔIKK^: Heterozygous Tmem119^Cre^ mice were bred to be homozygous Ikbkb^fl/fl^. Experimental animals were generated by crossing Tmem119^Cre/+^::Ikbkb^fl/fl^ mice with Ikbkb^fl/fl^ mice. Ikbkb^fl/fl^ littermates were used as controls. Primers were designed using NCBI Primer3, see Table S1 for sequences.

### Induction of CreER-mediated recombination

Cre-ER-mediated recombination in the Tmem119^ERΔIKK^ model was induced using 5 subcutaneous injections of tamoxifen (Sigma T5648-1G) dissolved in purified corn oil (Sigma C8267) 24h apart based on body weight for a total concentration of 2mg/kg body weight.

### Data sources, inclusion criteria and meta-analysis

We conducted a search on PubMed to identify published studies that examine the association between environmental benzene exposure and the prevalence of metabolic diseases. Our search strategy involved utilizing specific keyword combinations, including (“volatile organic compound” OR “benzene exposure”) AND (“metabolic disease” OR “insulin resistance”) in the abstract or title of the articles. Articles were excluded if they lacked metabolic measurements, were primarily methodological in nature, or utilized animal models. The studies meeting our rigorous selection criteria included assessments of urinary t,t-MA levels alongside adjusted odds ratios (OR) for the risk of metabolic diseases, incorporating key parameters like blood glucose, insulin, and cholesterol levels. We conducted a meta-analysis using the fixed-effect model to estimate the pooled odds ratio (OR) and its associated 95% confidence interval (CI) for the association between benzene exposure and insulin resistance (HOMA-IR). The fixed-effect model operates on the assumption that all included studies share a common true effect size, with observed variation primarily attributed to random error. We performed this analysis using the statistical software R, version 4.2.3. To evaluate heterogeneity among the selected studies, we employed the Chi-square test (Cochran’s Q), with statistical significance defined as a p-value less than 0.1. The I² statistic was also used to quantify the proportion of total variation in effect size estimates attributable to heterogeneity. An I² value exceeding 50% was considered indicative of substantial heterogeneity among the studies. Heterogeneity: Chi-square = 83.60, p <0.001, *I*^2^= 91.63%.

### Benzene exposure

The mice in inhalation chambers using FlexStream™ automated Perm Tube System (KIN-TEK Analytical, Inc) were exposed to benzene concentration of 50 ppm for 6h/day (acute) or up to 4 weeks (chronic) as described before ^18^. FlexStream™ automated Perm Tube System allows creating precision gas mixtures. This unit provides a temperature-controlled permeation tube oven, dilution flow controls, and front panel touch-screen interface. Mixtures are produced by diluting the miniscule flow emitted from Trace Source™ permeation tubes with a much larger flow of inert matrix gas, typically nitrogen or zero air. Control animals were breathing filtered air.

### Energy Balance

In order to assess VO_2_ consumption, VCO_2_ production, respiratory exchange ratio (RER) and heat production mice were placed in PhenoMaster metabolic cages (PhenoMaster, TSE system, Germany #160407-03). Animals were individually housed during 12 hours dark cycle (6 pm – 6 am). The mice were acclimatized for 48h and data was collected for 72h while food and water were provided *ad libitum*.

### Continuous glucose monitoring (CGM) system

To monitor glycemic changes, we used continuous glucose monitoring (CGM) system. CGM provides blood glucose readings automatically every 15 minutes. The commercial Libre 2 glucose monitor device (Abbott Diabetes Care), was engaged to obtain the assembled glucose sensor. After mice were anesthetized by inhalation with 2.5% Isoflurane, the sensor was inserted using a 23-gauge needle to pierce the shaved caudal surface of the skin. Subsequently, the sensor was scanned periodically using the Reader to obtain the continuous glucose monitoring (CGM) data. Animals were allowed 3 days for acclimatization after CGM implantation.

### Glucose tolerance test

For glucose tolerance test (GTT), mice were fasted for 6 hr (05:00-11:00) and intraperitoneally injected with D-glucose at a dose of 2 g/kg⋅BW. Blood glucose levels were measured at basal state (0 min) and then at 15, 30, 60, 90, and 120 minutes after injection as before ^90^. Blood glucose levels were measured at the indicated times via tail vein bleeding using OneTouch glucometer. Blood insulin was determined on serum from tail vein bleeds using Insulin ELISA kit (Crystal Chem. Inc.).

### Perfusion and Histology

Mice were anesthetized (I.P.) with avertin and transcardially perfused with phosphate-buffered saline (PBS) (pH 7.5) followed by 4% paraformaldehyde (PFA). Brains were post-fixed, sank in 30% sucrose, frozen in OCT medium and then sectioned coronally (30 µm) using a Leica 3050S cryostat. Four series were collected and stored at –80°C in cryo protectant until processed for immunohistochemistry as previously described ^91^. For immunohistochemistry, free-floating brain sections were washed in PBS, blocked using 3% normal donkey serum (NDS) and .3% Triton X-100 in PBS and then stained with a primary antibody overnight in blocking buffer. For pMAPK, immunostaining, sections were pretreated for 20 min in 0.5% NaOH and 0.5% H_2_O_2_ in PBS, followed by immersion in 0.3% glycine for 10 min and placed in 0.03% SDS for 10 min then stained with primary pMAPK (1:100, anti-rabbit, Cell Signaling Technologies,#ab9102) antibody overnight. Other floating brain sections were washed with PBS several times; blocked for 1h in 0.3% Triton X-100 with 3% normal donkey serum in PBS; and then stained with the following primary antibodies overnight at 4⁰C: FoxO1 (1:200, anti-rabbit, Cell Signaling Technologies, #ab2880), TNFα (1:100, anti-rabbit, Cell Signaling Technologies, #ab11948), GFAP (1:500, anti-chicken, Millipore, #AB5541), Iba1 (1:1000 anti-goat, Wako, #ab5076), GFP (1:1000 anti-chicken, Abcam, #ab13970). All floating brain sections were washed with PBS several times; and incubated with the anti-rabbit, anti-chicken, anti-gout Alexa Fluor 488, 568, and/or 647 (Invitrogen, 1:200) secondary antibody for 2h. Sections were mounted onto Superfrost Plus slides (Fisher Scientific) and coverslips added with ProLong Anti-fade mounting medium (Invitrogen). Images were visualized with Nikon 800 fluorescent microscope using Nikon imaging DS-R12 color cooled SCMOS, version 5.00. For high-resolution, images were taken using a multiphoton laser-scanning microscope (LSM 800, ZEISS) equipped with a 63X objective.

### Morphology assessment

Immunofluorescence images were taken using a confocal laser-scanning microscope (LSM 800, ZEISS) equipped with a 63X objective for the arcuate nucleus (ARC) and ventromedial nucleus (VMH) of the hypothalamus. Stacks of consecutive images taken at 0.35 μm intervals were sequentially acquired, and 30 optical sectioning, along the optical axis (z-axis), were reconstructed to 3D using Fiji-Image J. The maximum and minimum thresholds were set up equally for all the images. The skeleton analysis of astrocyte and microglia branch length and endpoint were perfumed as the protocol as described by (Young and Morrison 2018), using Fiji-Image J. All microscopy images and quantifications were performed by investigators that were blinded to the sample’s ID.

### Microglia isolation and purity

Animals were perfused using 5-7 ml of cold DPBS and collected to tubes with DPBS. Each brain was cut into sagittal sections and placed into C-tubes. Digestion enzymes from the neural tissue dissociation kit were added according to the manufacturer instructions (#130-092-628, Miltenyi Biotec) and homogenized using gentleMACS^TM^ dissociator system (#130-093-235, Miltenyi Biotec). Cell suspension was passed through a pre-wet 70 µm strainer. Following the debris removal (#130-109-398, Miltenyi Biotec), red blood cell lysis (#130-094-183, Miltenyi Biotec) and dead cell removal (#130-090-101, Miltenyi Biotec) live cells were incubated with 10ul of the CD11b (microglia)-Microbeads (#130-093-634, Miltenyi Biotec) for 15 min at 4 °C. After the incubation excess CD11b-Microbeads were washed out by centrifuging at 300 x g for 10 min with 1-2 ml of FACS buffer. Cells were resuspended in 500 ul FACS buffer (2% Fetal bovine serum and 1x PBS) and passed through the pre-wet MS separation columns (#130-042-201, Miltenyi Biotec) attached to the OctoMACS^TM^ Separator (#130-042-109, Miltenyi Biotec). MS separation columns were washed three times using 500 ul FACS buffer and flushed out with 1 ml of FACS buffer. For better purity flushed out cell suspension was passed again in new MS separation column. 50 ul of CD11b^+^ microglia fraction was tested on the Nothern light instrument for microglia purity and the remaining CD11b^+^ microglia fraction was processed immediately for the RNA extraction. Purity of microgila was assessed by full-spectrum flow cytometry (Northern Lights, Cytek Biosciences, Fremont, CA). Briefly, debris and cell aggregates were excluded based on laser scatter, dead cells were excluded based on propidium iodide (PI) staining, and erythrocytes were excluded based on differential side scatter ^92^. Astrocytes were defined as ASCA-2+CD45- and, within the remaining non-astrocytes, microglia as CD11b+CD45dim.

### Microglia RNA extraction

CD11b^+^ microglia fraction was centrifuged at 300g for 10 min and pelleted cells were resuspended in 0.75 ml of TRIzol^TM^ (Invitrogen #A33251) in phase-maker tubes. Cells were incubated for 5 min to permit complete dissociation of the nucleoproteins complex. Following the addition of 0.2 ml of chloroform per 1 ml of TRIzol^TM^ and incubation for 2-3 minutes samples were centrifuged for 5 min at 12000-16000 x g at 4 °C. After separating the aqua phase into a new tube and adding 0.5 ml of isopropanol per 1 ml of TRIzol™, samples were incubated overnight at –80 °C, and later centrifuged for 10 min at 12000 x g at 4 °C. Supernatant was discard and 1 ml of 75 % ethanol per 1 ml of TRIzol^TM^ was added to the samples and centrifuged for 5 min at 7500 x g at 4 °C. Supernatant was discarded and resulting pellet was air dried for 5-10 min. The pellet was resuspended in 15 ul of RNase-free water. RNA was quantified with a Nanodrop OneC spectrophotometer (#ND-ONEC-W, ThermoFisher Scientific) and quality-assessed by HSRNA ScreenTape (#5067-5579, Agilent Technologies) with a 4150 TapeStation analyzer (#G2992AA, Agilent Technologies).

### Hypothalamic RNA extraction

Male mice were sacrificed at *ad libitum* to harvest the brain and to isolate the hypothalamus. Hypothalamus samples were lysed with 0.75 ml of 2-mercaptoethanol added lysis buffer (PureLink^®^ RNA Mini Kit #12183025). Following the homogenization, with 0.75 ml of 70% ethanol, samples was transferred to spin cartridges with the collection tubes. Samples were centrifuged at 12000 x g for 14 sec at room temperature. After washing the samples three times with washing buffer, cartridges were centrifuged at 12000 x g for 1-2 min to dry the membrane with bound RNA. RNA was eluted using 15-20 ul of RNase-free water and stored at –80 °C for future use.

### RNA sequencing and data analysis

Microglia library construction from 2 to 100 ng RNA and RNA-seq was performed by Genomic Sciences Core of the Oklahoma following protocols as described previously in detail (Ocañas et al., 2022). Briefly poly-adenylated RNA was captured, captured mRNA was eluted and cDNA libraries were prepared according to the manufactured instructions. Libraries were sequenced using an Illumina Novaseq 600 system (SP PE50bp, S4 PE150). CIBERSORTx, or digital cytometry was used to estimate the cell type abundance from bulk microglia transcriptomics ^93^. https://cibersortx.staford.edu/ is used to create a signature matrix ^93^. The RNAseq files were run along with the created signature matrix to estimate the ratio of each cell type within our samples. CIBERSORTx generated results are shown as a heat map in Supplementary Figure 6. Hypothalamic RNA-Seq was performed at the WSU Genome Sciences Core. RNA concentration was determined by NanoDrop (ND-1000 UV-Vis Spectrophotometer) and quality was assessed using RNA ScreenTape on a 4200 TapeStation. RNA-seq libraries were prepared according to the QIAseq Stranded RNA Library Kits protocol before sequencing on a NovaSeq 6000 (2 x 50 bp). Microglia and hypothalamic RNA sequences were mapped to the mouse reference genome (GRCm38.90) using HISAT2 v.2.1.0.13 following the adapter trimming and quality checking. Quantification of the gene expression was generated using HTSeq-counts v0.6.0. Both microglia and hypothalamic RNA seq read counts data were analyzed using iDEP ^94^. The data was filtered to remove reads below .5 counts per million in at least one sample(s). The data was then transformed with EdgeR using a pseudo-count of 4, and missing values were inputted using the gene Median. Samples were visualized for using t-distributed stochastic neighbor embedding (tNSE) ^95^. Counts of RNA from individual samples were processed using DESeq2 to discover differentially expressed genes (DEGs) with FDR cutoff of <.1 and minimum fold change of >2. DEGs were further processed for pathway analysis using the Gene Set Enrichment Analysis (GSEA), for the enrichment of and gene ontology (GO) terms for biological process. DEGs were then referenced back to enriched pathways using GO categories and gene lists from the mouse Gene Ontology project at Mouse Genome Informatics (https://www.informatics.jax.org/) to create graphs of DEGs that appear in enriched pathways. Heat maps, bubble plots, and gene expression changes were plotted using SRplot (https://bioinformatics.com.cn/). All RNA-Seq data are available at the Sequence Read Archive (SRA) at NCBI under accession number PRJNA1035111.

### Serum corticosterone levels

The concentrations of corticosterone in blood were determined using a Mouse Corticosterone Competitive EIA ELISA Kit - LS-F67102; LifeSpan BioSciences, Inc.) according to manufacturer protocol.

### Statistical analysis

Statistical analyses for differentially expressed mRNAs were performed pairwise using EdgeR in the software R (3.2.2). For all other experiments, results are expressed as the mean ± standard error and were analyzed using Statistica software (version 10). Graphs were generated using GraphPad Prism software. Sample size is annotated within figure legends. An analysis of t-test was used when compared only benzene treatment versus control, variance (ANOVA) with repeated measurements was used to analyze CGM, VO_2_ consumption, VCO_2_ production, RER, heat production and GTT. Other parameters were analyzed by two-way ANOVA. All data were further analyzed with Newman-Keuls post hoc analysis. The level of significance (α) was set at 5%.

## Notes

### Competing Interest Statement

The authors have declared no competing interest.

