## Supplementary data for "Microglial NF-κB Signaling Deficiency Protects Against Metabolic Disruptions Caused by Volatile Organic Compound via Modulating the Hypothalamic Transcriptome"

### SUPPLEMENTAL INFORMATION

**Figure S1: Effects of benzene exposure on glucose levels, corticosterone, and long-term impact on energy homeostasis, related to Figure 2.** Transdermal sensor for continuous glucose monitoring (CGM) was implanted and following 3 days for acclimatization animals were exposed for 6 hours during the light cycle from 9 AM to 3 PM. (A) day 1; (B) day 2; (C) day 3; (D) day 5; (E) day 6; (F) day 7; Energy homeostasis parameters measured during the dark cycle 7 days after the 4-week exposure in male mice (G) Respiratory exchange ratio (RER); (H) Heat production [kcal/h/;kg]; (I) Corticosterone blood levels assessed at 24 h, 4 days and 4 weeks (5d, 6h/d) following benzene exposure. Error bars show SEM for n = 8-10 mice/group. Repetitive way ANOVA with the Newman-Keuls post hoc test (*$p<0.05$, **$p<0.01$, ***$p<0.001$).

**Figure S2: Female mice exposed to benzene do not exhibit impaired energy homeostasis, related to Figure 2.** Energy homeostasis parameters following a 4-week exposure (5d, 6h/d) in female mice. (A) Oxygen consumption ($VO_2$) consumption [ml/h/kg]; (B) Carbon dioxide production ($VCO_2$) [ml/h/kg]; (C) Respiratory exchange ratio (RER); (D) Heat production [kcal/h/;kg] measured during the dark cycle. Error bars show SEM for n = 8-10 mice/group. All values are presented as mean ± SEM.

**Figure S3: Impaired hypothalamic insulin signaling after acute benzene exposure, related to Figure 3.** (A) Representative immunostaining images of FoxO1 (red) and DAPI (blue) in the ventromedial nucleus of the hypothalamus (VMH) in 12-week-old male mice exposed to benzene for 6 hours, euthanized 15 minutes after insulin injection (i.p.; 3 I.U./kg B.W.) or saline. White arrows in the confocal images (63x) (B) indicate FoxO1 protein localization (C indicates cytoplasmic, N indicates nuclear). Scale bars: 200 μm and 10 μm in the merged picture; (C) Percentage of cytoplasmic FoxO1 expression in the VMH, with error bars representing SEM for n = 3-4 mice per group. (D) Representative immunostaining images of pMAPK (green) in the ARC and VMH in 12-week-old male mice exposed to benzene for 6 hours, euthanized 15 minutes after insulin injection (i.p.; 3 I.U./kg B.W.) or saline. Scale bar: 200 µm; Quantification of pMAPK$^{+}$ cells per field in the ARC (E) and VMH (F); error bars indicate SEM for n = 3-4 mice/group. Images were analyzed from at least three to four sections. Repetitive way ANOVA with the Newman-Keuls post hoc test (*p<0.05, **p<0.01, ***p<0.001).

**Figure S4: Rescue of benzene-induced astrogliosis and mitigation of hypothalamic insulin resistance in Cx3cr1$^{GFP\Delta IKK}$ mice.** (A) Representative images demonstrating GFAP staining (in blue) in the ARC of 12-week-old control (IKK$^{fl/fl}$) and Cx3cr1$^{GFP\Delta IKK}$ male mice. Scale bar: 10 µm; Quantification of GFAP$^{+}$ astrocytes in the ARC (B) and VMH (C) (grey, IKK$^{fl/fl}$; green, Cx3cr1$^{GFP\Delta IKK}$); (D) Skeleton images representing astrocytes morphology in the ARC of 12-week-old control (IKK$^{fl/fl}$) and Cx3cr1$^{GFP\Delta IKK}$ male mice; (E) Astrocytes endpoints (µm); (F) Astrocytes branch length (µm) (grey, IKK$^{fl/fl}$; green, Cx3cr1$^{GFP\Delta IKK}$); (G) Representative immunostaining images of FoxO1 (red) and DAPI (blue) in the ARC and (I) VMH. Scale bars: 200 µm. Percentage of cytoplasmic FoxO1 expression in the ARC (H) and VMH (J); (K) Representative immunostaining images of pMAPK (green) in the ARC in 12-week-old Cx3cr1$^{GFP\Delta IKK}$ male mice exposed to filtered air or benzene for 6 hours, euthanized 15 minutes after a single dose of insulin injection (i.p.; 3 I.U./kg B.W.) or sterile saline (light green, Cx3cr1$^{GFP\Delta IKK}$ saline-treated; dark green, Cx3cr1$^{GFP\Delta IKK}$ insulin-treated). Scale bar: 200 µm; (L) Quantification of pMAPK$^{+}$ cells per field in the ARC; error bars indicate SEM for n = 3-4 mice/group. Images were analyzed from at least three to four sections. Repetitive way ANOVA with the Newman-Keuls post hoc test (*p<0.05, **p<0.01, ***p<0.001).

**Figure S5: Validation of microglia purity in adult mice isolated from brains after 24-hour exposure to benzene.** (A) Flow cytometry of the CD11b-MACS positive microglia; (B)

CIBERSORTx analysis for microglia, astrocytes, neurons and oligodendrocytes cells in filtered air and benzene exposed samples.

**Figure S6: Benzene exposure does not affect body weight or fasted blood glucose in TMEM119$^{ER\Delta IKK}$ male mice.** (A) Body weight (g); (B) Fasted blood glucose (mg/dl) of 12-week-old control (IKK$^{fl/fl}$) and TMEM119$^{ER\Delta IKK}$ male mice exposure to filtered air or benzene for 4 weeks; error bars indicate SEM for n = 5-12 mice/group.

**Table S1. Genotyping primers, related to STAR Methods.**

| Genotyping Primers | Source | Identifier |
|---|---|---|
| Primers for Ikbkb flox allele;<br>*Forward*: GATCCGGAACCCTTAATATAACTTC<br>*Reverse*: CAACAAACTGACCACATAGCTCAAT | | N/A |
| Primers for Cx3cr1(GFP) allele;<br>*Forward*: TTAATGACCTGCAGCCAAGCTA<br>*Reverse*: TGTATGCTATACGAAGTTATTAGGTCTGAAGA | | N/A |
| Primers for Cx3cr1(KI WT) allele;<br>*Forward*: GCATATTCTTCATCACCGTCATCAG<br>*Reverse*: GGCGGCCAGGACGAT | | N/A |
| Primers for TMEM119 allele;<br>*Common reverse*: CGGGTCCCAGCAGTTCT<br>TMEM119 KO<br>*Forward*: GGGAGGCAGAGGGTTTCC<br>TMEM119 WT<br>*Forward*: TCCCTGTGCCTGCAACA | | N/A |

S1

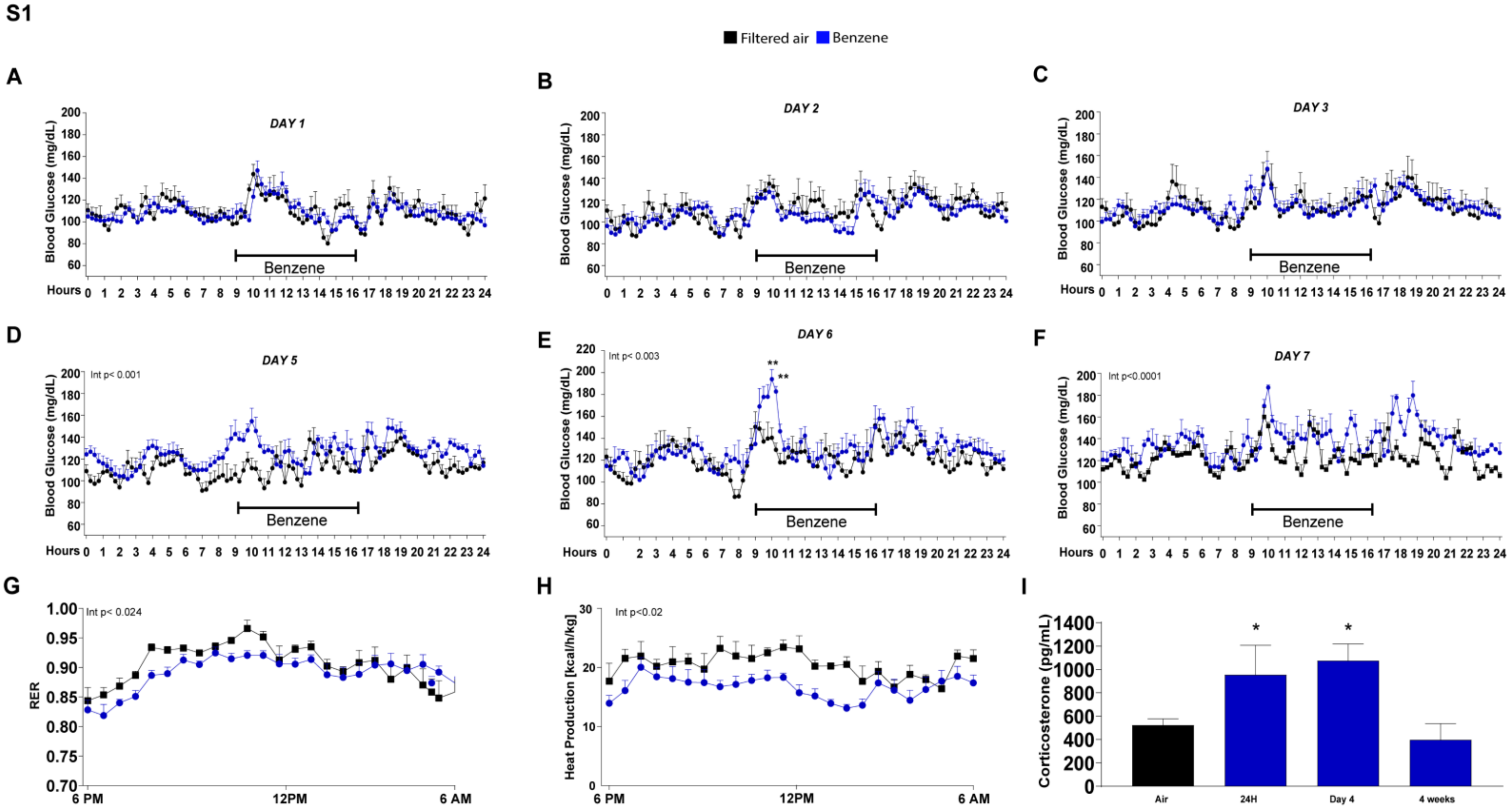
Filtered air Benzene
A
DAY 1
Blood Glucose (mg/dL)
Benzene
Hours
B
DAY 2
C
DAY 3
D
DAY 5
Int p< 0.001
E
DAY 6
Int p< 0.003
F
DAY 7
Int p<0.0001
G
RER
Int p< 0.024
6 PM
12PM
6 AM
H
Heat Production [kcal/h/kg]
Int p<0.02
I
Corticosterone (pg/mL)
Air
24H
Day 4
4 weeks

**Figure S1**

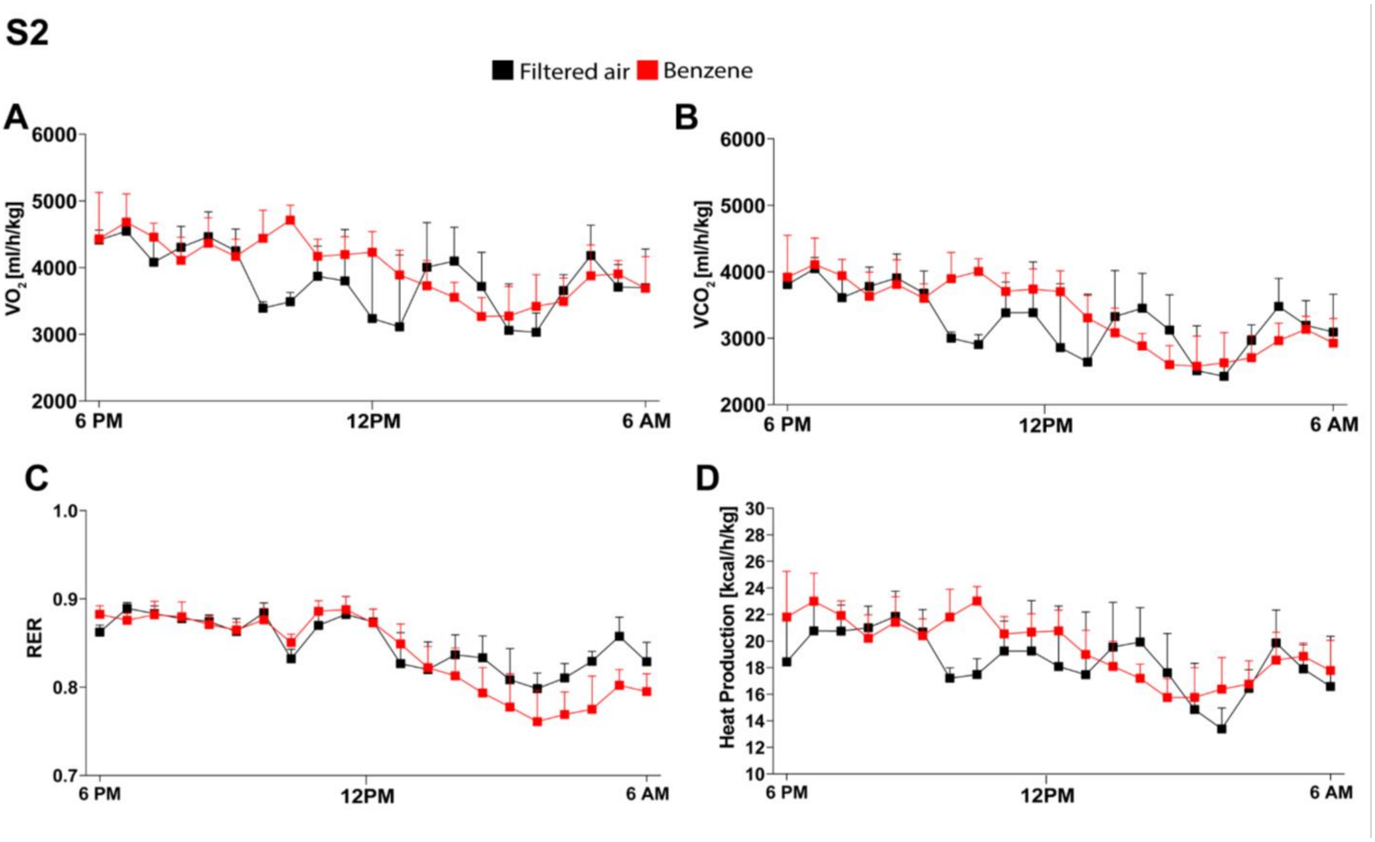
S2
Filtered air
Benzene
A
VO2 [ml/h/kg]
6000
5000
4000
3000
2000
6 PM
12PM
6 AM
B
VCO2 [ml/h/kg]
6000
5000
4000
3000
2000
6 PM
12PM
6 AM
C
RER
1.0
0.9
0.8
0.7
6 PM
12PM
6 AM
D
Heat Production [kcal/h/kg]
30
28
26
24
22
20
18
16
14
12
10
6 PM
12PM
6 AM

**Figure S2**

S3

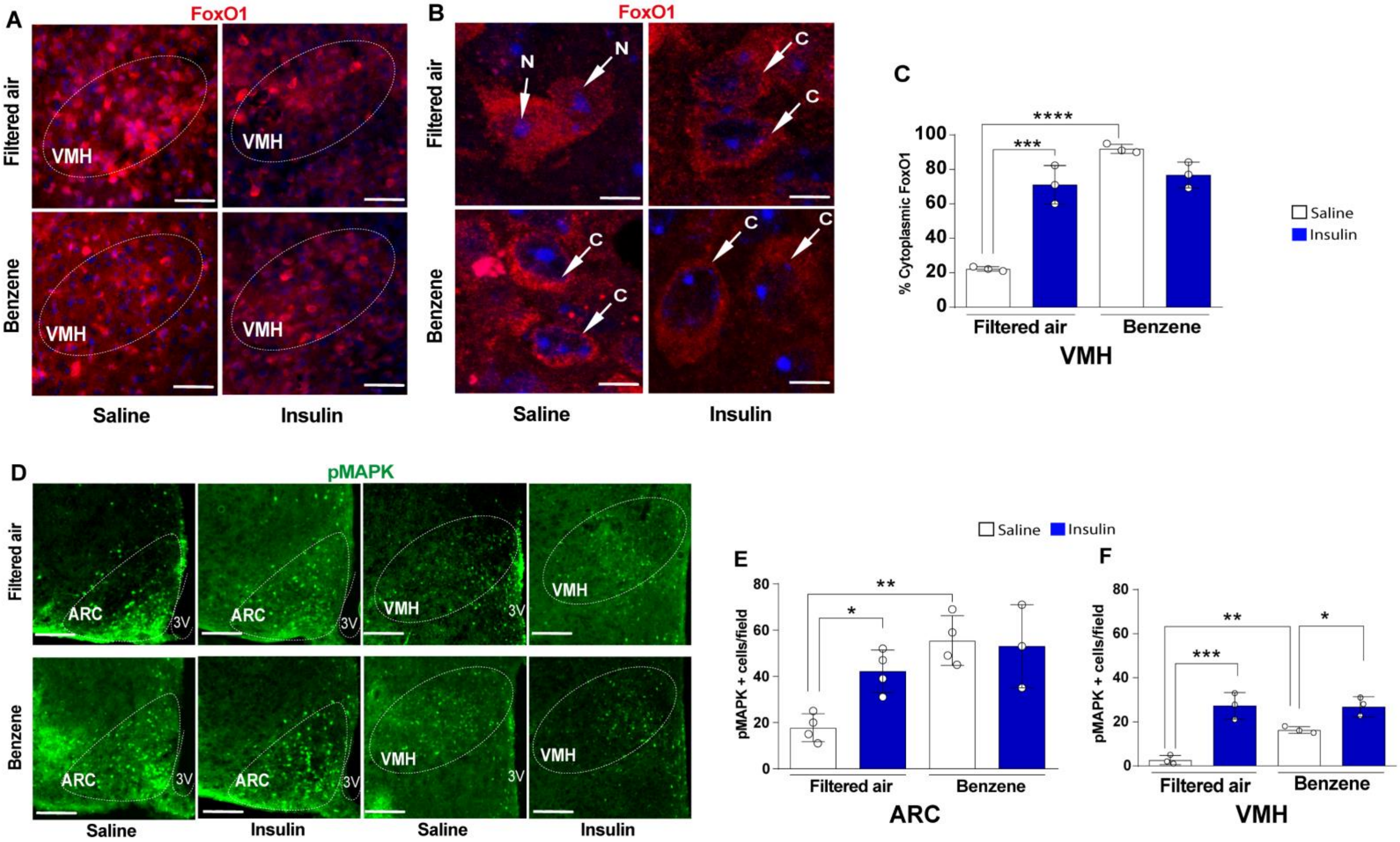
A
FoxO1
Filtered air
Benzene
VMH
Saline
Insulin
B
FoxO1
N
C
C
% Cytoplasmic FoxO1
100
80
60
40
20
0
****
***
Filtered air
Benzene
VMH
Saline
Insulin
D
pMAPK
ARC
3V
E
F
pMAPK + cells/field
**
*
***
ARC

**Figure S3**

S4

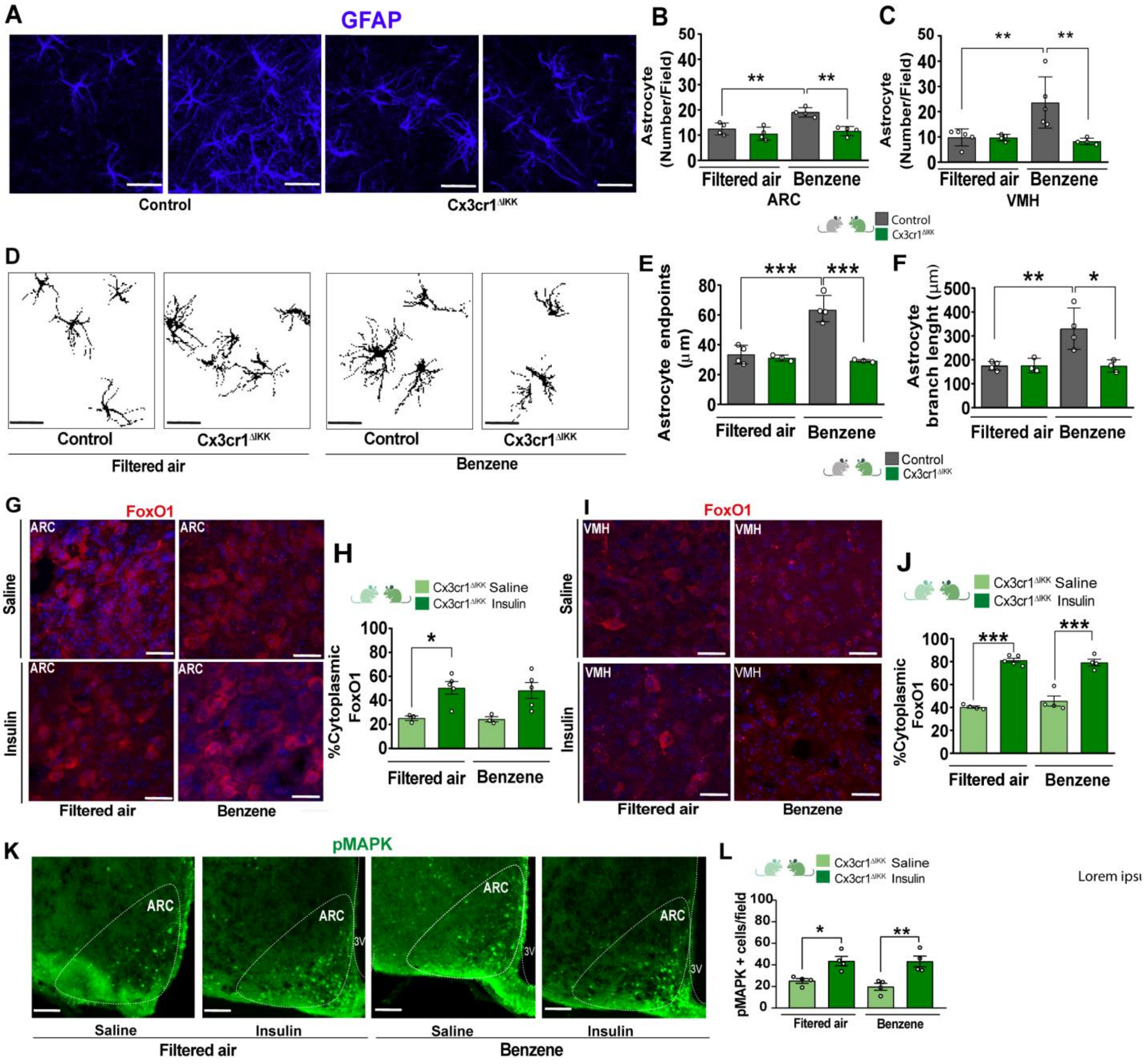
A
GFAP
Control
Cx3cr1^ΔIKK
B
Astrocyte (Number/Field)
**
**
Filtered air
Benzene
ARC
C
Astrocyte (Number/Field)
**
**
Filtered air
Benzene
VMH
Control
Cx3cr1^ΔIKK
D
Control
Cx3cr1^ΔIKK
Control
Cx3cr1^ΔIKK
Filtered air
Benzene
E
Astrocyte endpoints (μm)
***
***
Filtered air
Benzene
F
Astrocyte branch lenght (μm)
**
*
Filtered air
Benzene
Control
Cx3cr1^ΔIKK
G
FoxO1
ARC
Saline
Insulin
Filtered air
Benzene
H
Cx3cr1^ΔIKK Saline
Cx3cr1^ΔIKK Insulin
%Cytoplasmic FoxO1
*
Filtered air
Benzene
I
FoxO1
VMH
Saline
Insulin
Filtered air
Benzene
J
Cx3cr1^ΔIKK Saline
Cx3cr1^ΔIKK Insulin
%Cytoplasmic FoxO1
***
***
Filtered air
Benzene
K
pMAPK
ARC
3V
Saline
Insulin
Saline
Insulin
Filtered air
Benzene
L
Cx3cr1^ΔIKK Saline
Cx3cr1^ΔIKK Insulin
pMAPK + cells/field
*
**
Fitered air
Benzene
Lorem ipsı

**S5**

**A**

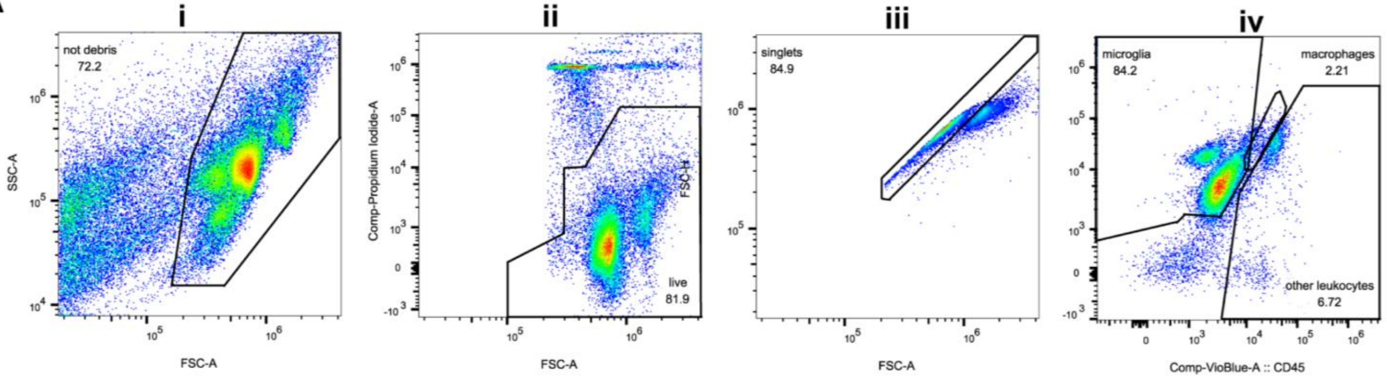


**B**

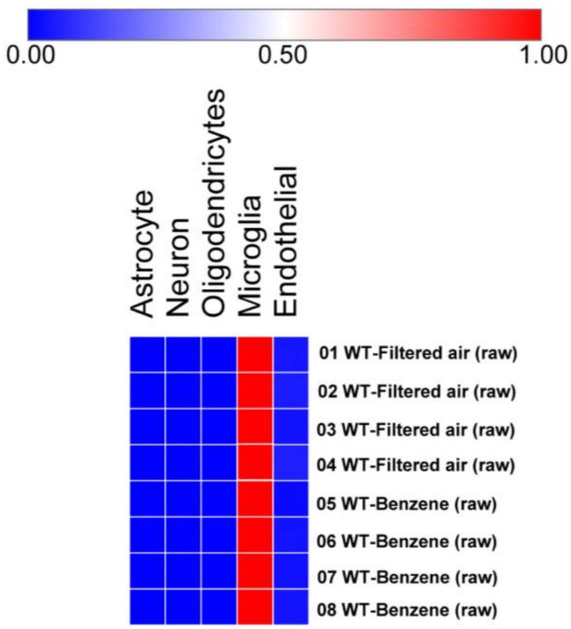


**Figure S5**

S6

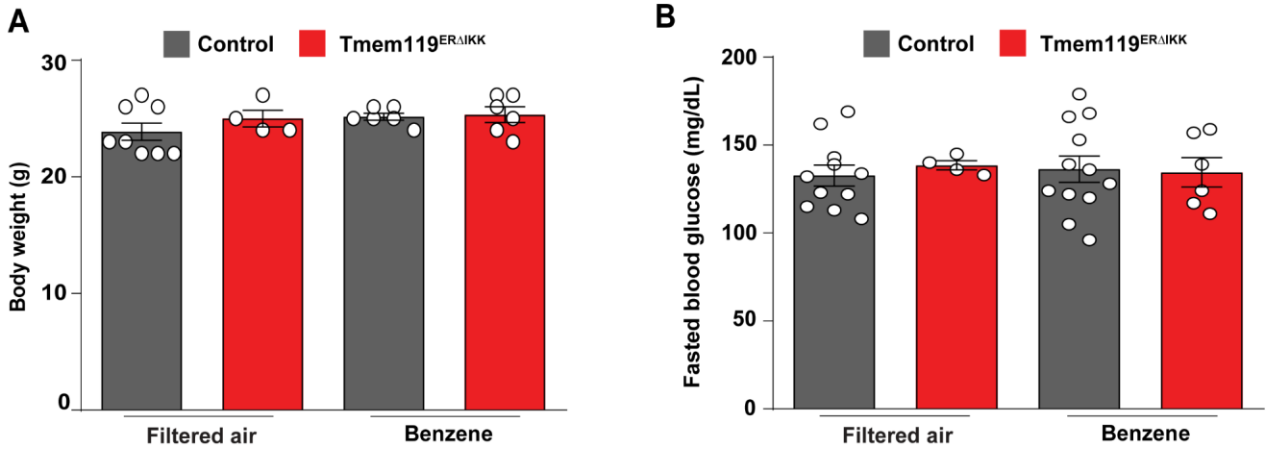
A
Control
Tmem119ERΔIKK
Body weight (g)
30
20
10
0
Filtered air
Benzene
B
Control
Tmem119ERΔIKK
Fasted blood glucose (mg/dL)
200
150
100
50
0
Filtered air
Benzene

**Figure S6**
